# Target complementarity in cnidarians supports a common origin for animal and plant microRNAs

**DOI:** 10.1101/2023.01.08.523153

**Authors:** Yael Admoni, Arie Fridrich, Talya Razin, Miguel Salinas-Saavedra, Michal Rabani, Uri Frank, Yehu Moran

**Affiliations:** Department of Ecology, Evolution and Behavior, Alexander Silberman Institute of Life Sciences, Faculty of Science, Hebrew University of Jerusalem, Jerusalem 9190401, Israel; Department of Genetics, Alexander Silberman Institute of Life Sciences, Faculty of Science, Hebrew University of Jerusalem, Jerusalem 9190401, Israel; Center for Chromosome Biology, School of Biological and Chemical Sciences, University of Galway, Galway, Ireland

**Author notes:** Correspondence: Yael Admoni, Yehu Moran **Email:**.

**Keywords:** MicroRNA, Cnidaria, *Nematostella*

## Abstract

microRNAs (miRNAs) are important post-transcriptional regulators that activate silencing mechanisms by annealing to mRNA transcripts. While plant miRNAs match their targets with nearly-full complementarity leading to mRNA cleavage, miRNAs in most animals require only a short sequence called ‘seed’ to inhibit target translation. Recent findings showed that miRNAs in cnidarians, early-branching metazoans, act similarly to plant miRNAs, by exhibiting full complementarity and target cleavage; however, it remained unknown if seed-based regulation was possible in cnidarians. Here, we investigate the miRNA-target complementarity requirements for miRNA activity in the cnidarian *Nematostella vectensis*. We show that bilaterian-like complementarity of seed-only or seed and supplementary 3’ matches are insufficient for miRNA-mediated knockdown. Furthermore, miRNA-target mismatches in the cleavage site decrease knockdown efficiency. Finally, miRNA silencing of a target with three seed binding sites in the 3’ untranslated region that mimics typical miRNA targeting was repressed in zebrafish but not in *Nematostella* and *Hydractinia symbiolongicarpus*. Altogether, these results unravel striking similarities between plant and cnidarian miRNAs consolidating the evidence for common evolutionary origin of miRNAs in plants and animals.

## Introduction

miRNAs are endogenous post-transcriptional regulators that are abundant in diverse eukaryotic lineages (Ameres & Zamore, 2013; Bartel, 2004, 2018; Moran *et al*, 2017). They have important roles in various biological processes and are essential for proper development of animals and plants (Bartel, 2018; Jones-Rhoades *et al*, 2006; Voinnet, 2009; Dexheimer & Cochella, 2020). miRNAs are transcribed by RNA polymerase II into long primary transcripts that are processed into hairpin-structured precursor-miRNAs (pre-miRNAs), which are later cleaved into short 20-22 nucleotides long duplexes. The duplexes are loaded into Argonaute (AGO) proteins that are a part of the RNA-Induced Silencing Complex (RISC), where only one strand is selected to remain loaded and the other is discarded (Kim *et al*, 2009). The loaded strand leads the RISC complex to a matching target and mediates its repression by inducing cleavage, translation inhibition or degradation by deadenylation of the mRNA (Jones-Rhoades *et al*, 2006; Jonas & Izaurralde, 2015).

The miRNA system is present in both plant and animal kingdoms, although a few major differences exist between them in the miRNA biogenesis pathway, mode of action and target recognition (Axtell *et al*, 2011; Moran *et al*, 2017). The biogenesis pathway in animals starts within the nucleus with the processing of the primary miRNA (pri-miRNA) by the microprocessor complex composed of RNase type III Drosha and its partner protein Pasha (known as DGCR8 in vertebrates) (Han *et al*, 2004a). The resulting pre-miRNA is transported by Exportin 5 into the cytoplasm where it gets cleaved into the mature miRNA by the RNase type III Dicer with the help of partner double-stranded RNA binding proteins such as Loqs and TRBP (Förstemann *et al*, 2005; Jouravleva *et al*, 2022; Fareh *et al*, 2016; Redfern *et al*, 2013; Wilson *et al*, 2015). In plants, both pri-miRNA and pre-miRNA are processed within the nucleus by DICER-LIKE1 (DCL1) assisted by its partner protein Hyponastic Leaves1 (HYL1) (Voinnet, 2009; Han *et al*, 2004b). Another difference between plant and animal miRNAs resides in their target recognition mode. In bilaterian animals, which include most known animal groups such as arthropods, nematodes, and vertebrates, miRNAs bind their targets with a short 5’ sequence called the ‘seed’ that includes only seven nucleotides, at positions 2- 8 of the miRNA (Brennecke *et al*, 2005). Supplemental complementarity at the 3’ end, mostly of positions 13-16, occurs in some cases, but it is not as frequent and considered less crucial for target recognition (Bartel, 2009; Grimson *et al*, 2007). The contribution of the supplemental complementarity to target binding seems to change considerably between cases (Becker *et al*, 2019; Bertolet *et al*, 2019). Target recognition restricted to seed or mediated via a seed match and supplemental complementarity with mismatches at positions 10 and 11 often leads to translational inhibition and deadenylation of the mRNA, mitigated by the metazoan-specific GW182 protein family (called TNRC6 in vertebrates) (Hutvagner & Simard, 2008; Bartel, 2009; Iwakawa & Tomari, 2015). Contrastingly, plant miRNA target recognition and activity require nearly-full complementarity that frequently results in AGO-mediated target cleavage between positions 10-11 of the miRNA, known as the cleavage site. Translational inhibition can also occur, but it still requires nearly-full complementarity (Iwakawa & Tomari, 2013).

The above-mentioned differences led to the notion that the miRNA system evolved independently in plants and animals; however, recent studies have shown that the miRNA system in the model sea anemone *Nematostella* vectensis, as well as other cnidarian species, is more similar to plants than previously thought (Moran *et al*, 2014; Modepalli *et al*, 2018; Tripathi *et al*, 2022; Baumgarten *et al*, 2018; Li & Hui). Cnidaria, the sister group to Bilateria, diverged over 600 million years ago from the vast majority of animal clades, and is composed of sea anemones, corals, jellyfish, and hydroids (Erwin *et al*, 2011; Kayal *et al*, 2018). Cnidarians possess a miRNA system (Grimson *et al*, 2008), and share some highly conserved miRNAs, some of them are known to be crucial for cnidarian fitness and development (Praher *et al*, 2021; Modepalli *et al*, 2018; Fridrich *et al*, 2020, 2023). Interestingly, cnidarian miRNAs operate by binding their targets with nearly-full complementarity that leads to mRNA cleavage, in a similar manner to plant miRNAs (Moran *et al*, 2014). Furthermore, miRNAs in *Nematostella* and plants are methylated at the 3’ end, which is essential to prevent miRNA degradation (Modepalli *et al*, 2018). Importantly, it was shown recently that *Nematostella* Hyl1-Like a, a homolog to plant-specific HYL1, also takes part in miRNA biogenesis, which suggests that it likely took part in miRNA biogenesis before the separation of plants and animals (Tripathi *et al*, 2022). Yet, despite these striking similarities to the miRNA pathway of plants, the cnidarian miRNA system also exhibit clear metazoan-specific features such as Drosha and Pasha homologs and a GW182 homolog (Mauri *et al*, 2017; Moran *et al*, 2013). The similarities to the bilaterian miRNA pathway raise the question whether cnidarian miRNAs might be able to target mRNAs via more restricted interactions focused on the seed region like their bilaterian counterparts.

To answer this question, we characterized the complementarity requirements between miRNAs and their targets in *Nematostella*. Using a reporter, we tested the efficiency of different complementarity patterns in promoting gene knockdown, such as the bilaterian-like seed match and a cleavage site mutated sequence. We utilized *TBP::mCherry* transgenic sea anemones that ubiquitously express mCherry fluorescent protein (Admoni *et al*, 2020) and injected to their zygotes different miRNA mimics to compare their effect on the expression of the fluorescent reporter.

## Results and Discussion

### Bilaterian-like matches fail to repress gene expression in *Nematostella*

It was previously shown that injection of short hairpin RNAs (shRNAs) to *Nematostella* zygotes leads to efficient knockdown of chosen targets (He *et al*, 2018). For our study, we designed a miRNA mimic (mimiR) based on an endogenous miRNA template to resemble native *Nematostella* miRNA precursors and better predict their processing by Dicer and the strand selection by AGOs. We used the pre-miRNA sequence of Nve-miR-2022, a highly conserved miRNA among cnidarians (Moran *et al*, 2014; Praher *et al*, 2021), and changed the mature miRNA sequence to nearly fully match the mCherry transcript. The target site is located in the 3’ UTR, since the majority of canonical miRNA sites are found in this region (Bartel, 2018). To be able to test the effect of different complementarity patterns on knockdown efficiency, we first generated mimiR with nearly-full complementarity to mCherry transcript (except for position 19, see materials and methods), that was later altered to resemble bilaterian miRNA binding sites that are based on seed match or mismatched in the cleavage site (**Figure 1A**).

**Figure 1.**
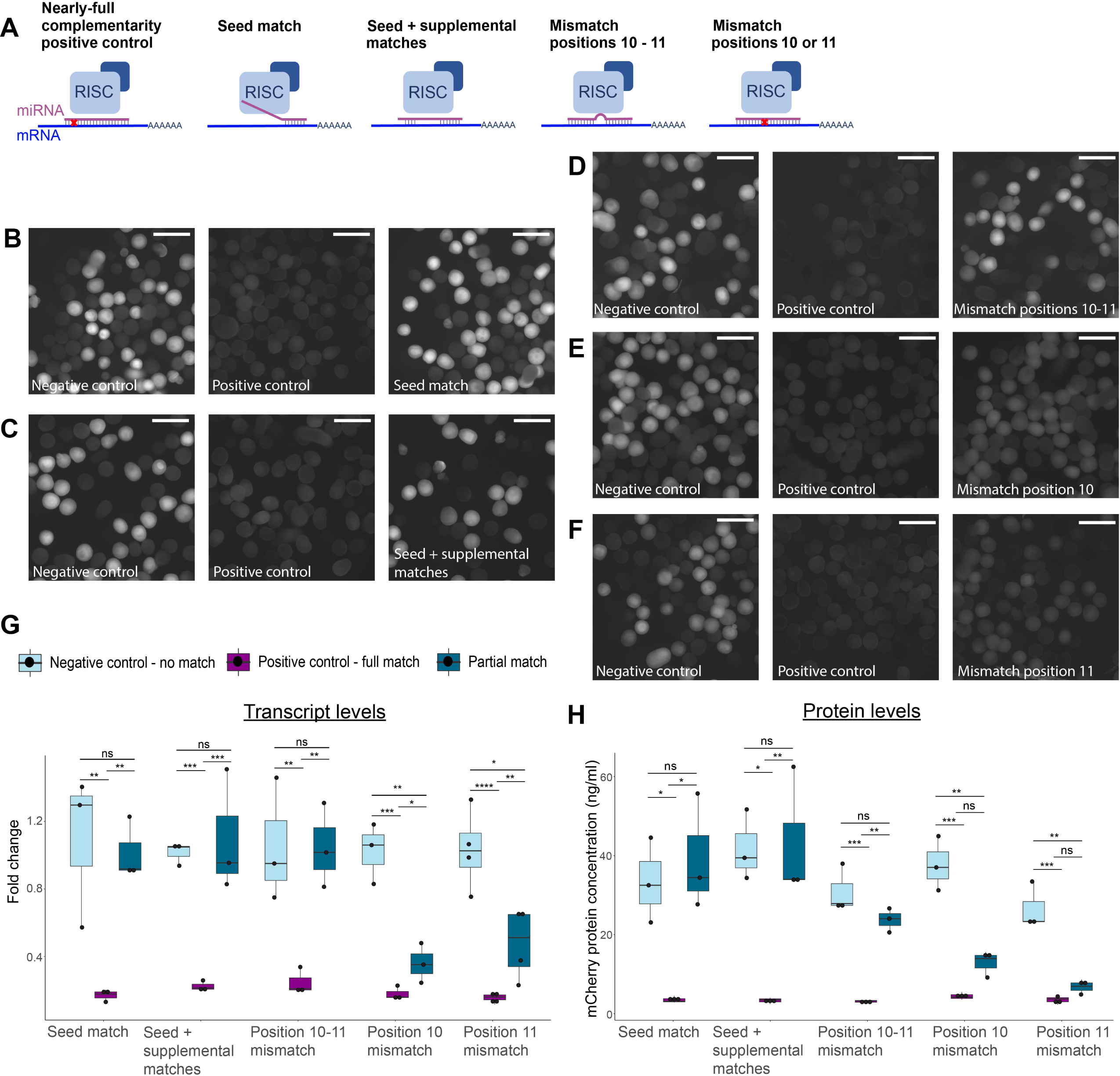
Silencing effects of different miRNA-target complementarity patterns in *Nematostella* A. Schematic representation of complementarity between designed mimiRs and mRNA target B-F. mCherry fluorescence captured in *TBP::mCherry* heterozygote embryos, three days after injection with mimiRs. Negative control groups (left) injected with shRNA with no match in *Nematostella* genome, displaying noticeable fluorescence. Positive control (middle) groups injected with nearly-full complementarity mimiR displaying no visible mCherry fluorescence. Groups injected with partial match mimiRs (right) displaying varying levels of fluorescence with (B) seed match (C) seed + supplemental matches and (D) mismatched positions 10-11 showing fluorescence similar to negative control and (E) mismatched position 10 and (F) mismatched position 11 showing intermediate fluorescence. Scale bars represent 500 µm. G. mCherry transcript fold change three days after injection with different mimiRs. Significance is shown for pairwise comparisons (one-way ANOVA with Tukey’s HSD post-hoc test, n=3 biological replicates for all groups but mismatched position 11 that has n=4 biological replicates). H. mCherry protein concentration three days after injection with different mimiRs. Significance is shown for pairwise comparisons (one-way ANOVA with Tukey’s HSD post-hoc test, n=3 biological replicates).

To test the impact of bilaterian-like miRNA-target complementarity pattern on transcript and protein levels, only the seed region at positions 2-8 of the miRNA was left matching the mCherry-encoding transcript while the rest of the sequence was changed. In addition, we generated a mimiR with supplemental matches to the seed at positions 13-16 (**Figure 1A**). The mimiRs were injected to *TBP::mCherry* zygotes that were observed after three days and mCherry transcript and protein levels were measured.

Interestingly, we observed that bilaterian-like mimiRs with ‘canonical site, i.e., matching their target only via the seed region, which is the most common class of miRNA in Bilateria (Bartel, 2018), cause no measurable knockdown of mCherry. The fluorescence of the embryos was similar to the negative control group injected with short hairpin RNA (shRNA) with no matches to *Nematostella* transcripts (**Figure 1B**) and mCherry mRNA and protein levels showed no difference compared to the negative control; and were significantly higher than in the positive control groups (**Figures G and H, and Table 1**). Adding supplementary binding bases at the 3’-end of the miRNA, to resemble another type of common bilaterian targets, resulted in similar measurements, with both transcript and protein levels of mCherry remaining unaffected by the presence of the mimiR when compared to the control group (**Figure 1C, G and H, and Table 1**).

**Table 1.**
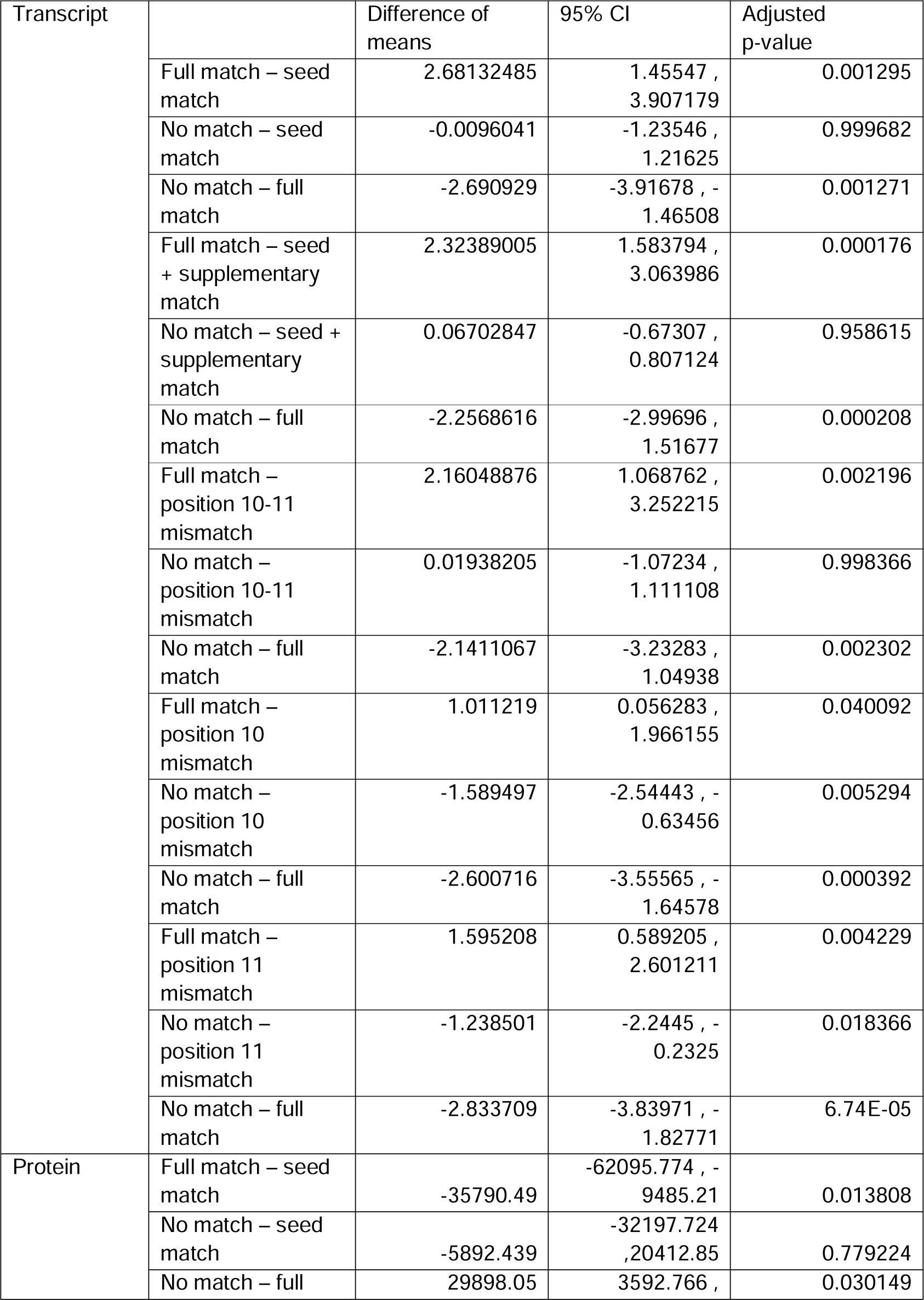
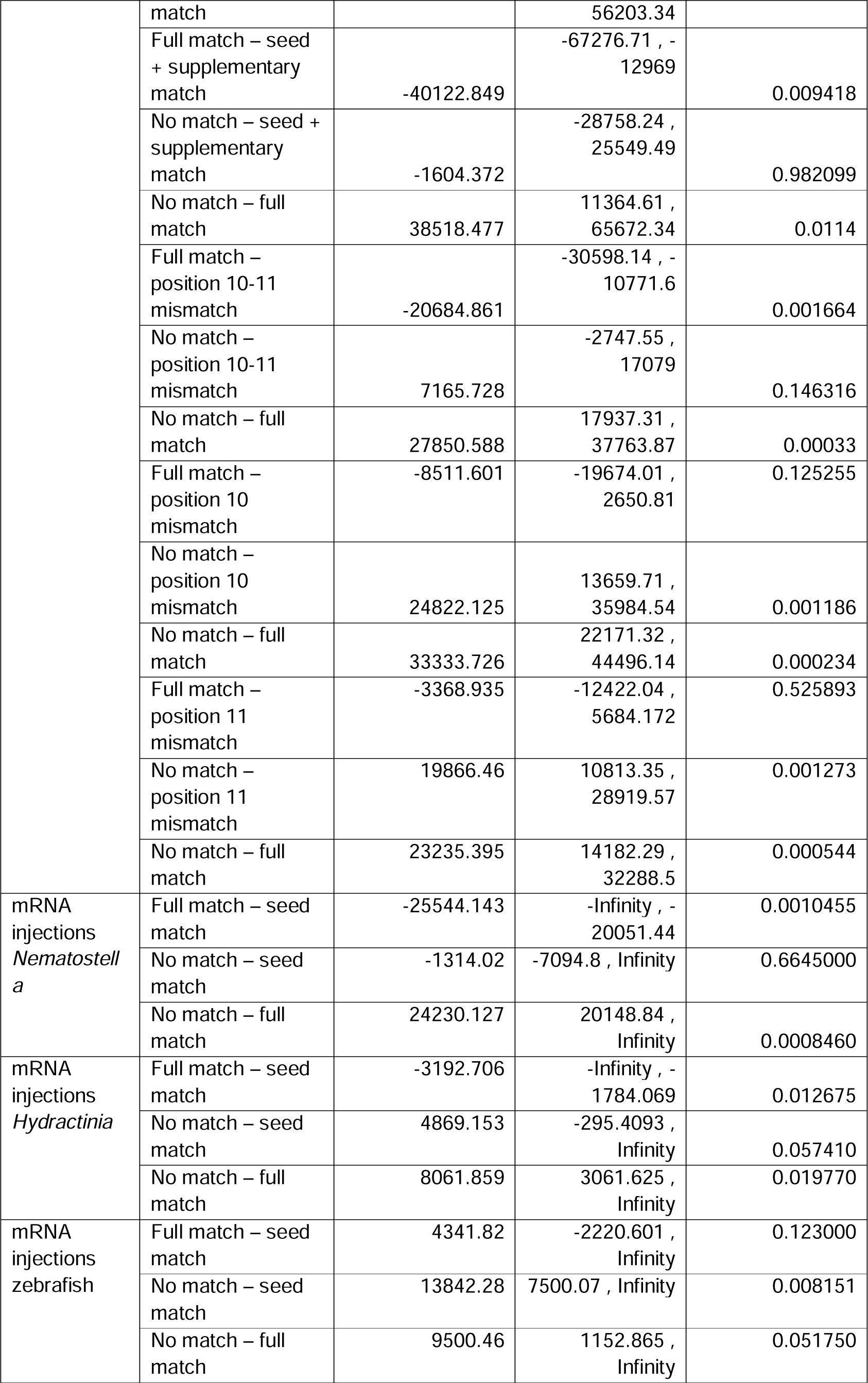
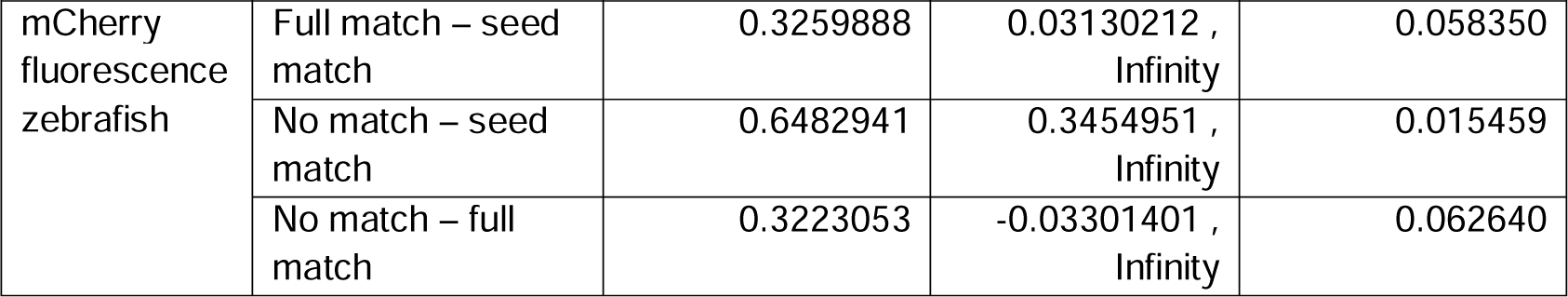
Pairwise comparisons. Results of pairwise comparisons between injected groups. Including difference of means for mCherry transcript (ΔCt), protein concentration or normalized fluorescence, confidence interval and adjusted p-value for multiple comparisons. P-values were adjusted with Tukey’s HSD for experiments with *TBP::mCherry* transgenic line and with FDR correction for mRNA injections. Full match refers to positive control group injected with nearly-full complementarity mimiR. No match refers to negative control group injected with shRNA with no match in *Nematostella* genome.

These results implicate that the previously described nearly-perfect plant-like matches between cnidarian miRNAs and their targets (Moran *et al*, 2014) are the major mode of interaction and bilaterian-like matches between miRNAs and targets are probably not functional in Cnidaria. In fact, it was shown that bilaterian-like matches have very weak effect or none at all in plants (Iwakawa & Tomari, 2013; Liu *et al*, 2014). Our experimental validation supports the evolutionary scenario that miRNA targeting based on seed match is a bilaterian innovation, suggested to contribute to the expansion of regulatory networks by allowing a single miRNA to bind hundreds of targets (Moran *et al*, 2017). It is noteworthy that in plants despite having full complementarity to the targets, the seed region is still crucial for target recognition and mismatches in positions 1-8 lead to decrease in silencing efficiency (Liu *et al*, 2014), which could potentially also be the case for *Nematostella* miRNAs.

### Cleavage site mismatches interfere with miRNA activity

Next, we assessed the necessity of miRNA binding in the site of target cleavage. We mismatched positions 10-11 of the mimiR and compared mCherry levels to nearly-full complementarity control mimiR injection. Similarly to seed-restricted mimiRs, inhibition of cleavage site binding resulted in impaired miRNA activity, as mCherry fluorescence as well as transcript and protein levels showed no difference from the negative control (**Figure 1D, G, and H, and Table 1**).

These results are in accordance with plant miRNAs that also fail to induce target cleavage when central mismatches are introduced (Iwakawa & Tomari, 2013). Moreover, this experiment further validates the notion that *Nematostella* miRNAs promote target cleavage as the main mode of action (Moran *et al*, 2014). Cleavage is known to be the main mechanism for miRNA activity in plants as well, suggesting that it is the ancestral miRNA mode of action. This has been discussed in relation to the ancient RNA interference (RNAi) system for defense against invasive nucleic acids, such as transposons and viruses, that operates by binding of short interfering RNAs (siRNAs) to foreign RNA targets and eliminating their expression by cleaving them. It is a probable evolutionary scenario that the miRNA system evolved from the RNAi defense system (Cerutti & Casas-Mollano, 2006).

In addition to target cleavage, translational inhibition, which is the common miRNA mechanism in bilaterians, was also exhibited in plants (Liu *et al*, 2014; Li *et al*, 2013; Brodersen *et al*, 2008). It was shown in *Arabidopsis thaliana* that central mismatches abolish target cleavage, although still allow translational inhibition when the target site is in the 5’ UTR (Iwakawa & Tomari, 2013). The extent to which translational inhibition is occurring in *Nematostella* is still unknown, although it was shown that *Nematostella* GW182 homolog could promote mRNA decay and translational repression when expressed in human cells (Mauri *et al*, 2017). Based on our experiments, it seems that translational inhibition or mRNA decay did not occur when central mismatches prevented target cleavage; however, we cannot exclude the scenario that this mechanism is active in *Nematostella* due to the conservation of GW182, but requires different complementarity pattern or different number of sites.

### Partial mismatch in the cleavage site results in weaker repression in *Nematostella*

After testing positions 10-11, we wished to test how a single mismatch in the cleavage site affects the knockdown efficiency. For this we mismatched either position 10 or 11 separately and injected both variants to transgenic zygotes. Both mimiRs resulted in visibly lower mCherry fluorescence. On the molecular level, mCherry transcript levels were significantly lower than the negative control, hence knockdown still occurred, but it was significantly weaker than the positive control inflicted repression (**Figure 1E-F and G, and Table 1**). On the protein level, despite a noticeable trend of higher protein levels compared to the positive control both in the ELISA measurement and the fluorescence of the transgenic animals, there was no statistically significant difference between protein concentration between a nearly-full match and a single position mismatch (**Figure 1E-F and H, and Table 1**). This raises the intriguing possibility that translational inhibition contributes to the silencing effect beyond the effect at the RNA level. Yet, such a claim requires further biochemical proof.

Reduced knockdown efficiency due to mismatch of one nucleotide at the cleavage site could be due to different conformation of AGO-miRNA that changes the cleavage efficiency (Sheu-Gruttadauria & MacRae, 2017). Some *Nematostella* miRNAs naturally exhibit a mismatch to their target in position 10 or 11: position 10 was found mismatched in 67 miRNA-target pairs, while position 11 was mismatched in 41 pairs among degradome-verified targets (Moran *et al*, 2014). It is possible that the natural mismatches are selected for weaker knockdown of their targets, as weaker repression might be beneficial for some regulatory roles.

### Multiple seed match sites in the 3’ UTR are inefficient for miRNA activity in Cnidaria

Single miRNA binding sites in bilaterians are capable of modulating protein target levels in a significant manner as demonstrated experimentally in *Drosophila* and zebrafish (Brennecke *et al*, 2005; Choi *et al*, 2007). However, many bilaterian miRNAs exhibit more than one site for each target they regulate (Grimson *et al*, 2007). Since multiple binding sites on the same target transcript can provide together synergic rather than additive repression effects (Briskin *et al*, 2020), we designed an mCherry-encoding mRNA that harbors three seed match sites in its 3’ UTR (**Figure 2A**). The mRNA was injected to wild-type *Nematostella* zygotes along with the previously used seed match mimiR. A new nearly fully matching positive control mimiR was designed since the original sequence could potentially partially bind the three seed sites.

**Figure 2.**
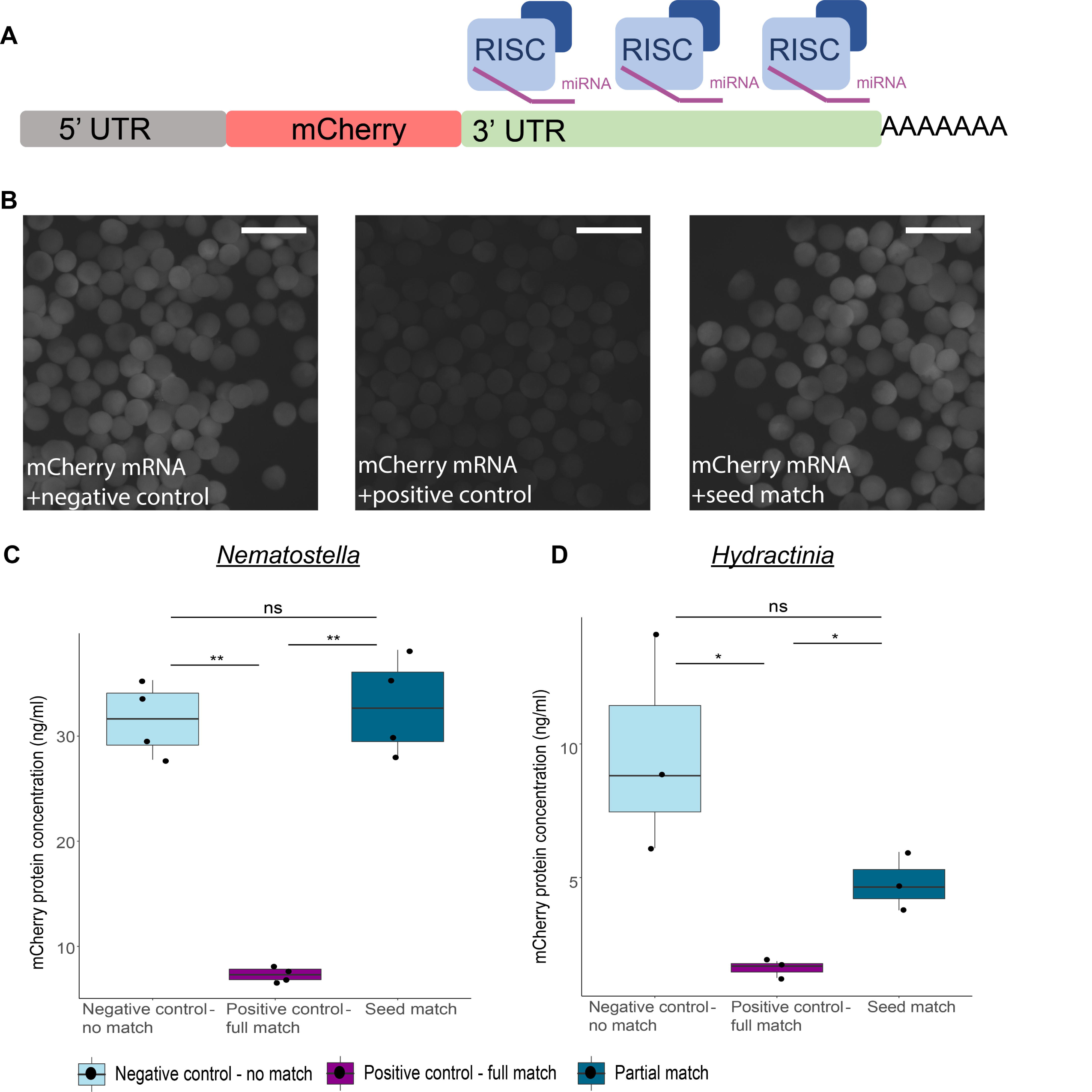
mRNA with multiple seed-match sites is not silenced by seed-match mimiR in *Nematostella* and *Hydractinia* A. Schematic representation of injected in-vitro transcribed mCherry mRNA, containing EF1α kozak sequence followed by mCherry-encoding transcript containing three seed-match sites in its 3’ UTR. B. mCherry fluorescence observed in WT *Nematostella* embryos, 24 hours after injection with mCherry mRNA combined with a different mimiR in each panel. Negative control group (left) injected with shRNA with no match in *Nematostella* genome, displaying noticeable fluorescence. Positive control (middle) group injected with nearly-full complementarity mimiR displaying no visible mCherry fluorescence. Seed match group (right) injected with mimiR matching the three seed sites in the 3’ UTR showing fluorescence similar to negative control. Scale bars represent 500 µm. C. mCherry protein concentration 24 hours after injection with mCherry mRNA combined with different mimiRs to *Nematostella* zygotes. Significance is shown for pairwise comparisons (One-tail Welch’s t-test with FDR correction, n=4 biological replicates). D. mCherry protein concentration 24 hours after injection with mCherry mRNA combined with different mimiRs to *Hydractinia* zygotes. Significance is shown for pairwise comparisons (One-tail Student’s t-test with FDR correction, n=3 biological replicates).

After 24 hours from injection, mCherry fluorescence was weaker in the positive control group compared to the seed match group (**Figure 2B**). In accordance, protein concentrations were similar between seed match mimiR and negative control, both significantly higher than the positive control (**Figure 2C Table 1**). This result shows that increasing the number of target sites in the 3’ UTR does not improve the efficiency of seed match miRNAs and further validates that bilaterian-like matches are ineffective in *Nematostella*.

Next, the mRNA silencing by mimiRs was tested in *Hydractinia symbiolongicarpus*, a colonial cnidarian and member of Medusozoa, which separated from Anthozoa, the group that includes *Nematostella*, about 560 million years ago (Khalturin *et al*, 2019). shRNA silencing tool was shown to be effective in *Hydractinia* (DuBuc *et al*, 2020), making it possible to test this experimental design in another cnidarian. As described before, mCherry protein levels were quantified 24 hours after injection to *Hydractinia* zygotes. Similar to *Nematostella*, the protein levels following injection of nearly-full complementarity mimiR were significantly lower than both the seed match and the negative control groups (**Figure 2D and Table 1**). In contrast, although there seems to be a slight difference between the negative control and the seed groups, it is not significant. These results further confirm that the miRNA mechanism in cnidarian animals is based on nearly-full complementarity between miRNAs and their targets, while seed match alone is insufficient in mediating target knockdown.

To validate the ability of the small RNAs to bind to the sites and promote silencing in a bilaterian system, the target mRNA was injected with shRNA/mimiR to zebrafish embryos. The injection was in the same manner as to *Nematostella*, with addition of an mRNA encoding sfGFP without miRNA binding sites, to account for the variability of expression efficiency in the zebrafish embryos. Following 10 hours from injection, the embryos show a difference in mCherry fluorescence between the groups, with the seed match group exhibiting the weakest fluorescence (**Figure 3A**). Both in the protein and the fluorescence level, a significant difference was found between the negative control and the seed match groups, hence validating the efficiency of the seed match mimiR in binding and repressing its target in a bilaterian animal (**Figure 3B and C and Table 1**). Moreover, no significant difference was found between the full match mimiR group and the other treatments, indicating that the target was not efficiently cleaved despite the extensive complementarity. This result is in accordance with the fact that zebrafish lack efficient slicing activity by AGO2 due to two specific point substitutions specific to teleost fishes (Chen *et al*, 2017). In conclusion, the results of this experiment validate the experimental approach of co-injection of seed match or nearly-full complementarity mimiR and the mRNA target since the results coincide with the literature about zebrafish miRNA pathway. Further, together with our previous results it suggests that seed-restricted matches between miRNAs and their targets is a derived bilaterian innovation (**Figure 4**).

**Figure 3.**
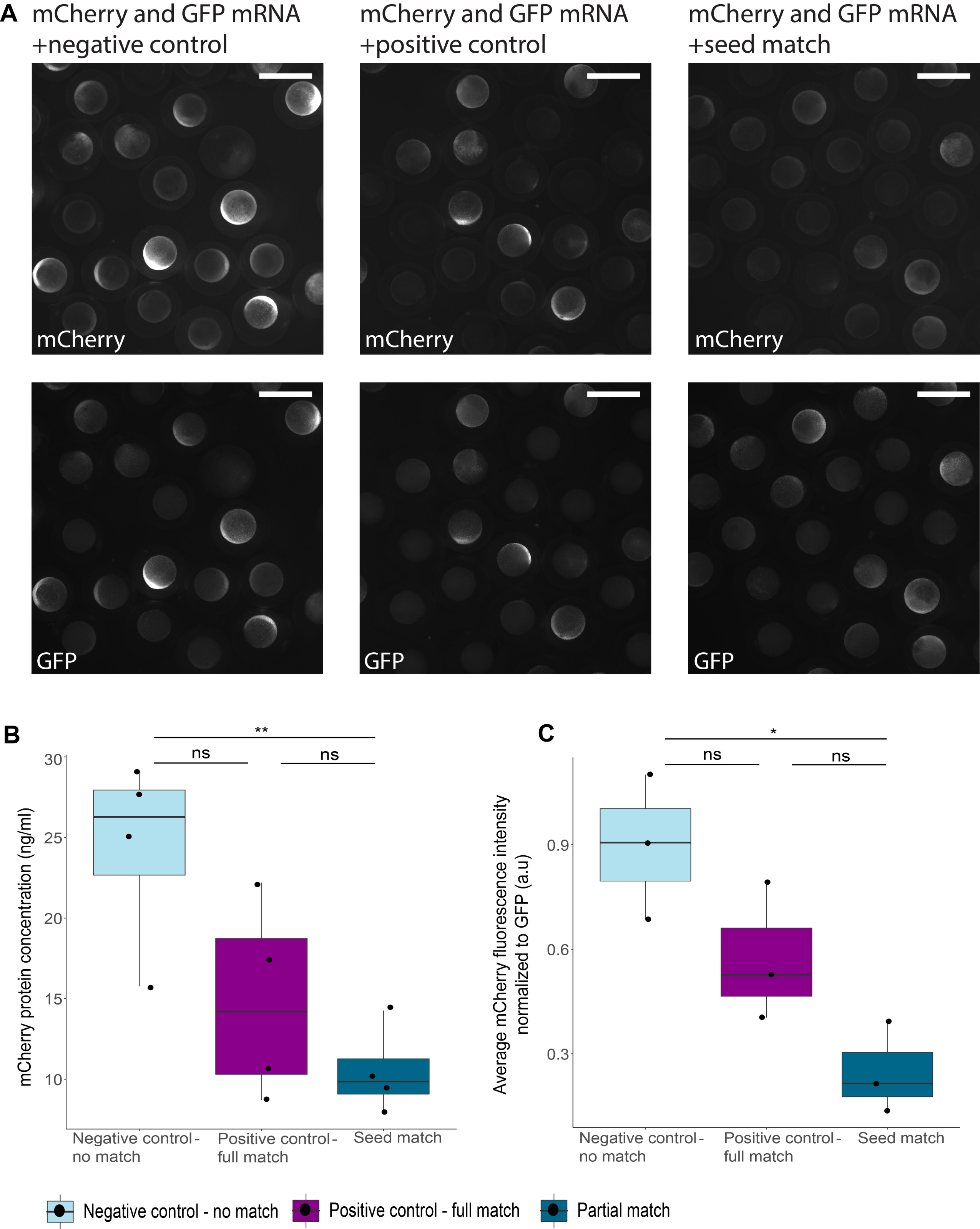
mRNA with multiple seed-match sites is repressed by seed-match mimiR in zebrafish A. Fluorescence observed in zebrafish embryos, 10 hours after injection with mCherry mRNA combined with different mimiRs and sfGFP mRNA as fluorescence intensity control. Top pictures show mCherry fluorescence in the different treatments: Negative control group (left) injected with shRNA with no match to mCherry mRNA, displaying noticeable fluorescence. Positive control (middle) group injected with nearly-full complementarity mimiR displaying weaker mCherry fluorescence. Seed match group (right) injected with mimiR matching the three seed sites in the 3’ UTR showing the weakest fluorescence of the groups. Bottom pictures show sfGFP fluorescence with variability in expression but don’t follow the same trend as mCherry fluorescence. Scale bars represent 1000 µm. B. mCherry protein concentration 10 hours after injection with mCherry mRNA combined with different mimiRs. Significance is shown for pairwise comparisons (One-tail Student’s t-test with FDR correction, n=4 biological replicates). C. Average fluorescence intensity of mCherry normalized to GFP 10 hours after injection with mCherry mRNA combined with different mimiRs and sfGFP mRNA as intensity control. Significance is shown for pairwise comparisons (One-tail Student’s t-test with FDR correction, n=3 biological replicates).

**Figure 4.**
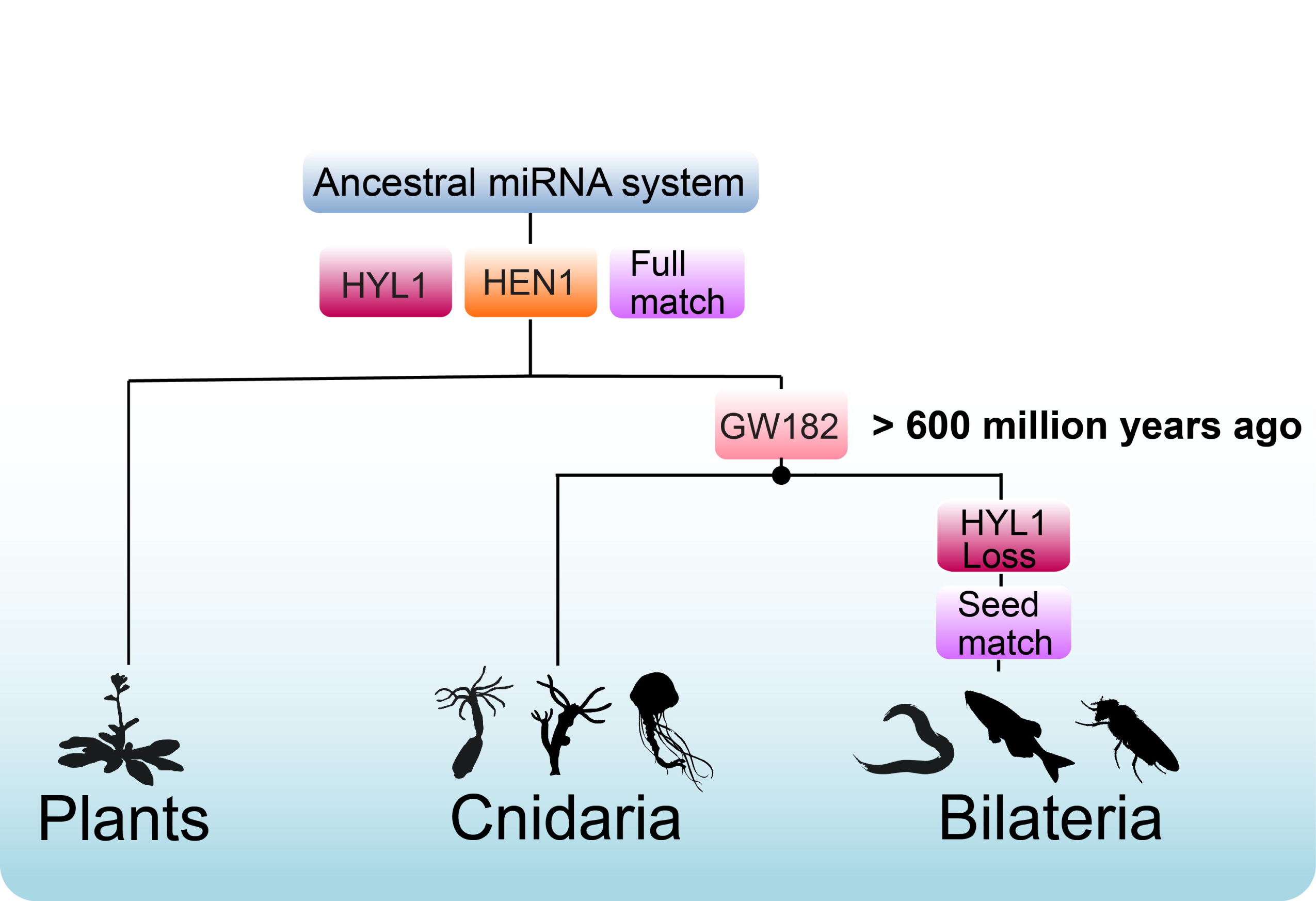
Evolution of the miRNA system in plants, Cnidaria and Bilateria Results presented here and in previous publications (Moran *et al*, 2014, 2017; Tripathi *et al*, 2022; Mauri *et al*, 2017; Modepalli *et al*, 2018) suggest that miRNA post-transcriptional regulation evolved before the separation of plants and animals. HYL1 and HEN1 were present in the common ancestor of plans and animals where they had roles in miRNA biogenesis and methylation of miRNAs to protect from degradation, respectively. GW182 was mitigating target translational inhibition before the separation of Cnidaria and Bilateria over 600 million years ago. While plant and cnidarian miRNAs match their targets with nearly-full complementarity, bilaterian miRNAs evolved to depend on seed-match. In addition, bilaterians lost HYL1 that was replaced with other miRNA biogenesis proteins. Animal silhouettes are from http://phylopic.org/.

In this study, we show through *in-vivo* assays that cnidarian miRNAs act similarly to those of plants in terms of the complementarity requirements to their targets to induce efficient gene repression. We show that bilaterian matches that rely either solely on seed matches or seed matches with supplementary 3’ matches fail to perform measurable gene repression in *Nematostella*. In addition, multiple seed matches are insufficient for promoting target silencing in *Hydractinia* and *Nematostella*. Furthermore, a cleavage site is crucial for miRNA activity in Cnidaria, and a single mismatch in the cleavage site reduces repression efficiency on the qualitative level.

Overall, the results of this study reveal important similarities between plants and cnidarians in the complementarity requirements between miRNAs and their targets and together with previous findings provide multiple lines of evidence for common origin of miRNA regulation before the separation of plants and animals (**Figure 4**).

## Materials and Methods

### *Nematostella* culture and microinjection

*Nematostella* polyps culturing, spawning and fertilization were conducted as previously described (Genikhovich & Technau, 2009) with minor modifications. Cultured anemones and were maintained at 18 °C under dark conditions and fed with freshly hatched *Artemia salina* nauplii three times a week. Anemones were induced to release gametes in 25 °C for 8 hours followed by fertilization of WT eggs with either WT or heterozygote *TBP::mCherry* sperm. The gelatinous sack surrounding the eggs was removed by incubation in 3% L-Cysteine (Merck Millipore, USA) while rotated by hand for 15 minutes. Microinjection to zygotes was performed with Eclipse Ti-S Inverted Research Microscopes (Nikon, Japan) connected to an Intensilight fiber fluorescent illumination system (Nikon) for visualization of the fluorescent injected mixture. The system is mounted with a NT88-V3 Micromanipulator Systems (Narishige, Japan). Every replicate included injection of three groups of 400-700 zygotes each: negative control shRNA group, positive control mimiR and altered mimiR. *TBP::mCherry* heterozygotes were injected with shRNA/mimiRs at 31.7 µM. WT zygotes were injected with mCherry mRNA at 0.167 µM along with shRNA/mimiR at 1 µM, and 100 mM KCl. All injection mixes included dextran Alexa Fluor (Thermo Fisher Scientific, USA) for tracing of injection mix. The injected animals were kept in an incubator at 22 °C, counted and transferred to fresh *Nematostella* medium (16 ‰ artificial sea water made from dry Red Sea salt) every day. The animals were visualized before flash-frozen in liquid nitrogen with ∼150 animals in each sample. The frozen samples were kept in -80 °C until either RNA or protein extraction.

### *Hydractinia* cultures and microinjection

Adult *Hydractinia symbiolongicarpus* colonies were maintained as previously described (Frank *et al*, 2020). The colonies were grown in artificial seawater at 19-22 °C on glass slides, separated to males and females. The animals were fed with *Artemia* nauplii four times per week, and once a week with ground oysters. The animals were kept in a constant 14:10 light:dark cycle, where females and males spawn 1.5 hours after exposure to light. Zygotes were injected within two hours from fertilization as previously described (Millane *et al*, 2011; Salinas-Saavedra *et al*, 2023), with mCherry mRNA at 0.167 µM and shRNA/mimiRs (Negative control shRNA, positive control mimiR or seed match mimiR) at 15.85 µM. Injected embryos were flash-frozen in liquid nitrogen after 24 hours. The frozen samples were kept in - 80 °C until protein extraction.

### Zebrafish embryos culture and microinjection

Wild-type zebrafish (AB/TL) maintenance was according to standard procedures. Fertilized eggs were collected at 28 °C and kept in culture medium (5 mM NaCl, 0.17 mM KCl, 0.33 mM CaCl_2_, 0.33 mM MgSO_4_, 0.25 mM HEPES, 0.1% Methylene blue). A total of ∼150 embryos per group were microinjected at the one-cell stage with 1 nl of solution containing 0.297 µM mCherry mRNA, 0.3 µM sfGFP mRNA and shRNA/mimiR at 1.782 µM (Negative control shRNA, positive control mimiR or seed match mimiR). All injection mixes included 10% phenol red (New England Biolabs) for tracing of injection mix. Zebrafish embryos were visualized under a fluorescent stereomicroscope before dechorionated and frozen at 10 hours post injection. For removal of chorion embryos were incubated for 5 mins with 1 mg/ml Pronase (Merck) then washed with culture medium and flash-frozen in liquid nitrogen. The frozen samples were kept in -80 °C until protein extraction. All protocols and procedures involving zebrafish were approved by the Institutional Committee on Animal Care and Use (IACUC, Protocol #NS-15859), The Alexander Silverman Institute of Life Sciences, The Hebrew University of Jerusalem.

### RNA extraction

Total RNA was extracted from ∼150 injected animals (3 days old planulae) with the aid of Tri-Reagent® (Merck Millipore) according to the manufacturer’s protocol, with a few minor changes. At the RNA isolation phase, samples were centrifuged at 21,130 *× g*. Removal of residual genomic DNA from the extracted RNA was conducted by treatment with Turbo DNase twice for 30 minutes at 37 °C (Thermo Fisher Scientific) and repeating the RNA purification procedure for a second time. Final RNA pellets were re-suspended in 23-25 µl of RNase-free water (Biological Industries, Israel). Final concentration was measured by Qubit™ RNA BR Assay Kit (Thermo Fisher Scientific. RNA integrity was assessed by gel electrophoresis with 1:1 formamide (Merck Millipore) and 1 µl of loading dye on 1.5% agarose gel. RNA samples were stored at -80 °C until used.

### shRNA /mimiR design

shRNA sequence to serve as negative control with no matches in *Nematostella* genome was taken from an existing protocol (Karabulut *et al*, 2019). mimiRs to target mCherry transcript were designed based on *Nematostella* miR-2022, an endogenous miRNA stem-loop that was used as template to allow better prediction of cleaving sites by Dicer and to ensure selection of the desired strand and loading onto AGO1 (Fridrich *et al*, 2020; Moran *et al*, 2014). The targeted sequence was selected have U as a 5’ terminal nucleotide, according to *Nematostella* guide strand characteristics, and mismatches were introduced to the predicted star strand in positions 1, 8, 9 and 17 (Fridrich *et al*, 2020). mimiRs were designed to the 3’ UTR region of the mCherry transcript. The base in position 19 was always cytosine, due to in-vitro transcription requirements. mimiR sequence alterations included mismatches in positions 10-11, position 10 or 11, only positions 1-8 base-pairing with the mCherry sequence (seed-match), and positions 1-8 and 13-16 matching (seed + supplemental matches).

### In-vitro transcription

The shRNA and mimiRs were transcribed according to the manufacturer’s instructions using the AmpliScribe™ T7-Flash™ Transcription kit protocol (Lucigen, USA) with a few changes. The shRNA/mimiR DNA templates were ordered from Integrated DNA Technologies, Inc (Integrated DNA Technologies, USA) as reverse complements to the sequence of T7 promoter followed by shRNA/mimiR precursor. Templates were annealed with a T7 promoter primer prior to the in-vitro transcription reaction, which was carried out for 15 hours, followed by addition of 1 µl of DNase, incubation at 37 °C for 15 min, and product clean up with Quick-RNA MiniPrep Kit (Zymo Research, USA). Concentration of transcripts was measured by Epoch Microplate Spectrophotometer (BioTek Instruments, Cole-Parmer, USA) and product integrity was validated with gel electrophoresis. The ready to use hairpins were kept at -80 °C until used.

### cDNA synthesis

cDNA synthesis was conducted with iScriptTM (Bio-Rad, USA) according to manufacturer’s protocol. 100 ng of RNA (extracted from ∼150 treated 3 days old planulae) per sample were used as template, resulting in final concentration of 5 ng/µl. cDNA was stored at -20°C.

### Reverse transcription-quantitative PCR

Primers to amplify mCherry transcript for RT-qPCR were designed via Primer3 version 0.4.0 (Untergasser *et al*, 2012) and calibrated at concentrations of 25, 5, 1, 0.2 and 0.04 ng/µl to generate standard curves with StepOnePlus Real-Time PCR System v2.2 (ABI, Thermo Fisher Scientific). Primer quality was 125% efficiency, -2.83 slope and R2 >0.99. The specificity of the amplified products was determined by the presence of a single peak in the melting curve. RT-qPCR was performed using StepOnePlus Real-Time PCR System v2.2 (ABI, Thermo Fisher Scientific) and cDNA amplification was quantitatively assessed with using Fast SYBR Green Master Mix (Thermo Fisher Scientific). Each sample was quantified in triplicates for mCherry transcript and housekeeping gene 4 (HKG4) as an internal control (Columbus-Shenkar *et al*, 2018). 1 µl of cDNA template was used for all replicates. For negative control cDNA was replaced with RNase-free water. The Reaction thermal profile was 95 °C for 20 sec, then 40 amplification cycles of 95 °C for 3 sec and 60 °C for 30 sec, a dissociation cycle of 95 °C for 15 sec and 60 °C for 1 min and then brought back to 95 °C for 15 sec (+0.6 °C steps). mCherry fold change was analyzed using a comparative Ct method (2^−ΔΔCt^) (Schmittgen & Livak, 2008). Thresholds for HKG and mCherry detection were equalized between individual experiments. Each experiment was composed of at least three biological replicates.

### Protein extraction

Total protein extraction was implemented by adding 200 µl of the following lysis buffer: 50 mM Tris-HCl (pH 7.4), 150 mM KCl, 10% glycerol, 0.5% NP-40, 5 mM EDTA (all chemicals purchased from Merck Millipore) and Halt™ Protease Inhibitor cocktail (Thermo Fisher Scientific). Then, samples were homogenized with pestle mixer (Argos Technologies, cat. No.A0001) and incubated at 4 °C for two hours in rotating mixer (Intelli Mixer™ RM-2, ELMI, function 1, 7 rpm). Samples were later centrifuged at 4 °C, 16,000 × *g* for 15 min and the aqueous phase was transferred to a new tube. Concentration of total protein was measured using Pierce™ BCA Protein Assay Kit (Thermo Fisher Scientific) on Epoch Microplate Spectrophotometer (BioTek Instruments). Samples were kept at -80 °C until used.

### Red Fluorescence Protein Enzyme-Linked Immunosorbent Assay (RFP ELISA)

mCherry protein levels were detected with the aid of RFP ELISA kit (Cell Biolabs, Inc., USA). All protein samples were diluted to equal concentration prior to loading on antibody plate, and the experiment was carried out according to protocol. Epoch Microplate Spectrophotometer (BioTek Instruments) was used for absorbance measuring. Fit for standard curve was found using CurveExpert Basic 2.2.3 (Hyams Development, USA).

### Rapid Amplification of cDNA Ends (3’ RACE)

In order to reveal the exact length of the 3’ end of the transgenic *TBP::mCherry* transcript, SMARTer® RACE 5’/3’ Kit was used (Takara Bio, Japan). Prior to the reaction, RNA was extracted from 3-months-old animals, with one round of Tri-Reagent® (Merck Millipore). cDNA synthesis was conducted according to manufacturer’s protocol with 500 ng of RNA. Gene specific primers were designed for the PCR and Nested PCR reactions, which were carried out by Advantage® 2 Polymerase (Takara Bio). Final products were outsourced for Sanger sequencing (HyLabs, Israel).

### mCherry mRNA generation

mCherry mRNA template was ordered as gBlock gene fragment (Integrated DNA Technologies). The sequence included T7 promoter, EF1α kozak sequence TGTTAAACCAACCAACCACC and 3’ UTR with three seed sites 21 bases apart (two were inserted in addition to the original one). In addition, the 3’ UTR included one site for full match mimiR and two nucleotides’ changes to make gBlock synthesis efficient. A codon-optimized mCherry mRNA sequence was design for expression in *Hydractinia*. The DNA fragment was dissolved in TE buffer to final 20 ng/µl and incubated at 50 °C for 20 min. For injection to *Nematostella*, the template was cloned to pGEM®-T Easy plasmid (Promega) and amplified with forward primer to add T7 promoter class II phi2.5. In-vitro transcription was conducted with HighYield T7 Cap 1 AG (3‘-OMe) mRNA Synthesis Kit (m5CTP) (Jena Bioscience, Germany) using 800 ng of amplified template followed by Turbo DNase treatment (Thermo Fisher Scientific) by incubation with 1 µl of DNase for 30 minutes at 37 °C twice sequentially. For injection to *Hydractinia* and zebrafish the mRNA template was amplified from gBlock, zebrafish mRNA with 68 °C annealing temperature. mRNA was transcribed with HiScribe T7 mRNA Kit with CleanCap Reagent AG (New England Biolabs) according to manufacturer’s protocol with 1 µg of amplified template. In-vitro transcription products were cleaned using RNA clean & concentrator 25 (Zymo Research) and eluted with 33 µl RNase free water.

Concentration was measured using Qubit ™ RNA Broad Range Assay Kit with the Qubit Fluorometer (Thermo Fisher Scientific). Poly-adenylation followed using *Escherichia coli* Poly(A) Polymerase (New England Biolabs) for 30 minutes at 37 °C and products were further cleaned with RNA clean & concentrator 5 (Zymo Research) and eluted with 8-10 µl RNase free water. Single product was validated on 1.5% agarose gel after incubated at 95 °C for two minutes in thermo cycler with hot lid then brought to 22 °C and mixed with formamide (Merck millipore) in 1:3 ratio. The mRNA was stored in -80 °C until injected.

### sfGFP mRNA generation

sfGFP-encoding mRNA with a 40 nucleotides polyA tail was in-vitro transcribed from a plasmid encoding the construct under SP6 promoter. The plasmid was linearized by digestion with SapI restriction enzyme (New England Biolabs) for 1 hour in 37 °C. mRNA was synthesized using HiScribe SP6 RNA synthesis kit (New England Biolabs) according to the manufacture’s protocol. Resulting mRNA levels were quantified by nanodrop (NanoDrop Microvolume Spectrophotometer, Thermo Fisher Scientific) and mRNA length was validated by gel electrophoresis. The mRNA was stored in -80 °C until injected.

### Microscopy

Fluorescence of mCherry and sfGFP protein was detected by SMZ18 stereomicroscope (Nikon) connected to an Intensilight fiber illumination fluorescent system (Nikon). Images were captured by DS-Qi2 SLR camera (Nikon) and were analyzed and processed with NIS-Elements Imaging Software (Nikon). Zebrafish images were taken using both mCherry and GFP channels and analyzed using ImageJ software (Schindelin *et al*, 2012). Between 17-29 zebrafish embryos were selected per field with three biological replicates per treatment. For quantitative analysis, background intensity calculated by averaging the intensity of the same five locations in each field was subtracted before raw intensity was measured for each individual embryo. mCherry intensity was normalized to GFP by dividing the values. The average normalized mCherry intensity was calculated for each treatment in three biological replicates.

### Statistical analysis

Comparisons between groups of injected animals in transcript levels and protein concentration were tested with one-way ANOVA with Tukey’s HSD post-hoc test. Normality of the data was validated beforehand. Statistical analysis for protein level or normalized mCherry intensity following mRNA injections was conducted with pairwise one-tail Student’s t-test or one-tail Welch’s t-test without the assumption of homogeneity of variances. P-values were adjusted by FDR method. For normalized mCherry intensity, average intensities were compared between groups. For mCherry transcript level, ΔCt values were compared between groups. All experiments included at least three biological replicates and three technical replicates for RT-qPCR and two for ELISA. The tests were performed in Rstudio 2021.09.0.

## Acknowledgments

The authors thank Dr. Reuven Aharoni (The Hebrew University of Jerusalem) for technical assistance. This work was supported by European Research Council grants CNIDARIAMICRORNA 637456 and AntiViralEvo 863809 to YM.

